# A Genome-wide CRISPR Screen Reveals ZDHHC8-Dependent G_α_q Palmitoylation as a Key Regulator of GPCR Signaling

**DOI:** 10.1101/2025.08.06.668953

**Authors:** Muhammad Ahmad, Raghuvir Viswanatha, Ah-Ram Kim, Norbert Perrimon

## Abstract

G protein-coupled receptors (GPCRs) that couple to the Gαq signaling pathway control diverse physiological processes, yet the full complement of cellular regulators for this pathway remains unknown. Here, we report the first genome-wide CRISPR knockout screen targeting a Gαq-coupled GPCR signaling cascade. Using a *Drosophila* model of adipokinetic hormone receptor (AkhR) signaling, we identified *CG34449 (Zdhhc8)*, encoding a palmitoyl acyltransferase and its adapter protein CG5447, as a top hit required for robust Gαq-mediated GPCR signaling. We show that Zdhhc8 enhances GPCR signaling through palmitoylation of Gαq, which promotes its membrane localization and function. Loss of *Zdhhc8* markedly reduces palmitoylation of Gaq resulting in attenuation of AkhR/Gαq signaling and a reduction in receptor stability. Mechanistically, Zdhhc8 is necessary for palmitoylation of Gαq. These findings uncover Zdhhc8-dependent Gαq palmitoylation as a pivotal regulatory mechanism in GPCR signal transduction and highlight palmitoyl transferase as potential modulators of GPCR pathways.

## Introduction

GPCRs are a large family of cell-surface receptors (over 800 in humans) that convert diverse extracellular stimuli into intracellular signals, thereby governing processes from neurotransmission and hormone release to immune and sensory functions^1^. Given their ubiquitous physiological roles, GPCR dysfunction is linked to numerous diseases^2^, and GPCRs remain prime therapeutic targets. Indeed roughly one-third of all approved drugs modulate GPCR signaling^1, 3^. Canonical GPCR signaling involves coupling to heterotrimeric G proteins: agonist-bound receptors act as GTP-exchange factors for the Gα subunit, causing dissociation of Gα–GTP from the Gβγ dimer and activation of distinct effectors^4, 5^. Different Gα families trigger specific second messengers, for example, Gαq activates phospholipase Cβ (generating inositol trisphosphate and mobilizing Ca2+), whereas Gαs and Gαi/o respectively stimulate or inhibit adenylyl cyclase (modulating cAMP levels). While the mechanisms regulating Gαs and Gαi/o mediated pathways have been extensively characterized, the modulation of Gαq signaling remains comparatively less understood^2, 6^. Notably, unlike Gαi/o and Gαs, which have long-standing research tools such as pertussis and cholera toxins, the Gαq family lacked selective inhibitors until the recent discovery of cyclic depsipeptide antagonists^7, 8^. The introduction of these Gαq-specific inhibitors has only begun to enable fine dissection of Gαq function, and relatively few studies to date have probed Gαq-driven signaling in depth. This gap highlights the need to elucidate how Gαq activity is regulated within native signaling networks ^1, 2^.

*Drosophila melanogaster* is a powerful genetic model to address this question, given the conservation of GPCR signaling and metabolic physiology between flies and mammals. The organs and endocrine pathways controlling energy homeostasis are functionally analogous in flies and humans, making *Drosophila* a valuable system to study metabolic GPCR pathways. One such pathway is governed by the adipokinetic hormone receptor (AkhR), a Class A GPCR that plays a central role in energy metabolism^9^. Adipokinetic hormone (AKH) is the primary insect hormone that mobilizes stored energy (lipids and trehalose) during high demand and is often regarded as the functional equivalent of mammalian glucagon^9, 10^. Binding of AKH to AkhR triggers a signaling cascade in the fly fat body (analogous to mammalian adipose and liver tissues) that promotes lipolysis and glycemia, thereby sustaining energy homeostasis during stress and activity ^11, 12^. Genetic studies have underscored the importance of this receptor: flies lacking AkhR are unable to efficiently catabolize fat stores and develop an obese phenotype, revealing that AkhR is essential for normal lipid mobilization. Despite the clear physiological significance of AkhR signaling, the molecular components and regulatory mechanisms downstream of AkhR (particularly its coupling to the Gαq pathway) remain incompletely defined.

In this study, we set out to dissect the AkhR signaling network and to uncover modulators of its Gαq-mediated pathway. We performed an unbiased, genome-wide CRISPR–Cas9 knockout screen in *Drosophila* cells, enabling the systematic discovery of intracellular factors that transmit or regulate the Akh hormonal signal. The CRISPR screen yielded a short list of high-confidence hits, prominently featuring two uncharacterized genes: *CG34449* and *CG5447*. *CG34449* encodes a polytopic membrane protein that we identify as the *Drosophila* ortholog of *Zdhhc8*, a member of the DHHC family of palmitoyltransferases. The second gene, *CG5447*, encodes a small cofactor protein anchored to the Golgi. CG5447 is the *Drosophila* ortholog of Golgin A7 (mammalian GOLGA7), which is known to form a complex with certain DHHC palmitoyltransferases at the Golgi membrane^13^. The convergent identification of a DHHC enzyme and its putative partner protein as top hits suggested a role of protein palmitoylation as a critical node in AkhR signaling. Here, we confirm this role by demonstrating that CG34449 (ZDHHC8) and its associated protein CG5447 are necessary for palmitoylation of Gαq.

## Results

### AkhR elicits robust Gαq-coupled calcium signaling and cytoskeletal remodeling in S2R^+^ cells

To establish a functional platform for interrogating Gαq-coupled receptor signaling, we generated stable *Drosophila* S2R^+^ cell lines expressing either the Adipokinetic hormone receptor (AkhR) or the Capa receptor (CapaR), which both engage Gαq pathways. Western blotting confirmed robust over expression of VHH05-tagged receptor constructs (**Figure 1A**) which appear as a high molecular weight smear consistent with extensive glycosylation of the receptor proteins.

**Figure 1.**
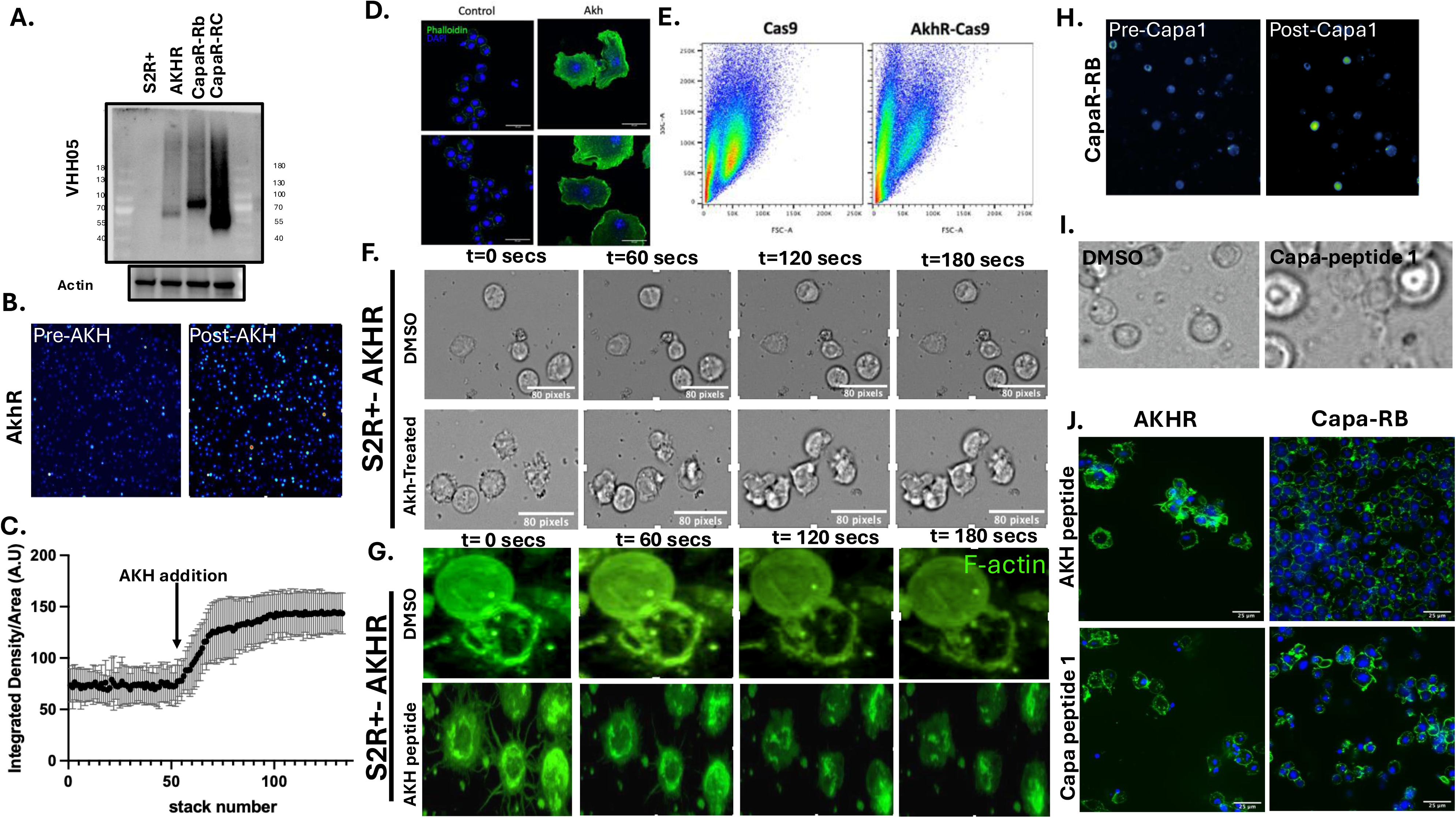
Distinct morphological and cytoskeletal responses to AkhR and CapaR activation in S2R⁺ cells. **(A)** Western blot confirming expression of VHH05 nano-tagged AkhR and CapaR in stably transfected S2R^+^ cell lines. **(B)** Calcium imaging using Fluo-4 AM shows increased cytosolic calcium upon AKH stimulation in AkhR-expressing cells. **(C)** Quantification of Fluo-4 fluorescence from (B) reveals a significant increase in intracellular calcium levels following AKH addition (arrow), indicating effective AkhR activation. **(D)** Confocal imaging of phalloidin-stained S2R^+^-AkhR cells reveals enhanced cell spreading after 48 hours of AKH treatment. **(E)** Flow cytometry analysis shows a rightward shift in side scatter (SSC) in AkhR-expressing cells following AKH stimulation, indicative of increased cell complexity; control Cas9-expressing cells do not show this shift. **(F)** Time-lapse brightfield imaging of S2R^+^-AkhR cells after AKH or DMSO addition. Membrane ruffling and dynamic morphological changes are observed within 180 seconds in AKH-treated cells but not in DMSO controls. **(G)** Live imaging of F-actin dynamics in S2R^+^-AkhR cells using live F-actin (cell mask) labeling confirms rapid cytoskeletal remodeling upon AKH stimulation. **(H)** CapaR-expressing cells exhibit increased cytosolic calcium after Capa-1 peptide stimulation, visualized by Fluo-4 AM staining. **(I)** Brightfield imaging of S2R^+^-CapaR cells reveals transient membrane protrusions following Capa-1 stimulation. **(J)** Comparative imaging of actin cytoskeletal responses in S2R^+^ cells expressing AkhR or CapaR following respective ligand stimulation. Phalloidin staining shows robust actin remodeling in AkhR-stimulated cells, while CapaR responses are comparatively weaker. Scale bars: 10 µm unless otherwise indicated.

Upon stimulation with synthetic AKH or Capa-1 peptides, both respective cell lines exhibited increased cytosolic calcium^14, 15^, as detected by Fluo-4 AM **(Supplementary Video 1).** However, AkhR-expressing cells consistently showed a stronger calcium response compared to CapaR (**Figure 1B–C**), indicating a more potent activation of the Gαq pathway by AkhR under these conditions.

We next investigated morphological changes downstream of receptor activation. Prolonged AKH treatment (48 h) of AkhR-expressing cells triggered pronounced cell spreading and actin remodeling, while Capa treatment of CapaR-expressing cells caused milder phenotypes (**Figure 1D, J**). Flow cytometry analysis revealed a substantial shift in side scatter (SSC) following AKH treatment of AkhR-expressing cells, consistent with cytoskeletal reorganization causing cell spreading. No comparable shift was observed in control (non-AkhR) cells (**Figure 1E**).

Live-cell imaging revealed that AKH stimulation triggered rapid membrane ruffling in AkhR-expressing cells treated with AKH peptide (**Supplementary Video 3**) compared to DMSO treated control (**Supplementary Video 2**), emerging within 60 seconds and intensifying over 3 minutes **(Figure 1F)**. Correspondingly, live F-actin imaging confirmed dynamic cytoskeletal remodeling in real-time **(Figure 1G).** Upon AKH peptide treatment, the actin cytoskeleton collapses inward and cells lose their surface attachments **(Supplementary Video 4)** in contrast to DMSO-treated controls (**Supplementary Video 3**). These effects were absent in DMSO treated control cells and were more robust than responses to Capa-1, as visualized by phalloidin staining (**Figure 1H–J**). These results establish AkhR as a robust Gαq-coupled receptor capable of driving rapid and sustained calcium mobilization^14^ and cytoskeletal remodeling in S2R^+^ cells, providing a sensitive system for probing downstream regulatory mechanisms. Based on this, we next selected clonal cell lines for VHH05-tagged AKHR and selected the highest expressing, clone 29 and Clone 39, for subsequent screening purpose **(Supplementary Figure 1A).**

### A genome-wide pooled CRISPR-Cas9 screen identifies regulators of AkhR-induced G**α**q signaling and cytotoxicity

Cytoskeletal changes are often accompanied by failures in cell division, a process that requires stereotypical cytoskeletal changes. Therefore, we reasoned that we could use cell differential cell enrichment in response to AKH peptide as a readout in a genome-wide CRISPR screen to identify AkhR signaling components. To test this hypothesis, we mixed AkhR-expressing cells transfected with either a GFP-expressing library vector or a BFP-expressing Gαq-targeting sgRNA. Upon AKH treatment, GFP^+^ (signaling-competent) cells displayed typical spreading and ruffling, while Gαq-knockout (BFP^+^) cells remained rounded and viable **(Figure 2A)**. Over 72 hours, GFP^+^ cells were progressively depleted at the expense of BFP^+^ cells, confirming that loss of Gαq conferred resistance to AKH **(Figure 2B)**. Next, AkhR-expressing cells were transduced with a genome-wide sgRNA library and exposed to AKH for 7 days alongside the BFP-labeled Gαq control population. Once selective enrichment of BFP^+^ cells was observed, genomic DNA was extracted and sgRNAs were sequenced **(Figure 2C).**

**Figure 2.**
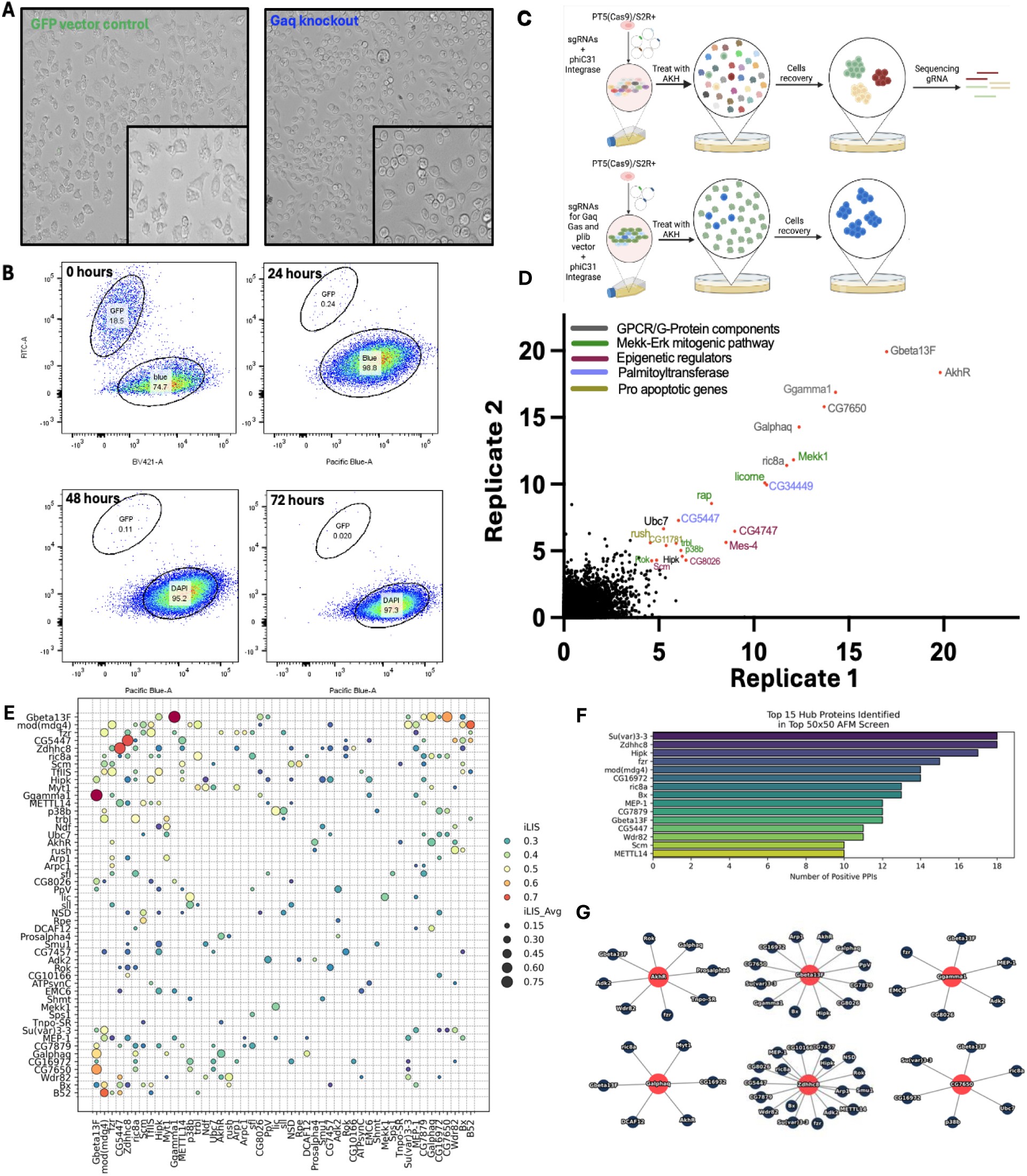
A pooled CRISPR-Cas9 screen identifies regulators of AkhR-mediated G_α_q signaling. **(A)** AkhR-expressing S2R+ cells were transfected with either a control GFP-expressing CRISPR library vector or a BFP-tagged construct encoding a sgRNA targeting G_α_q. Following treatment with AKH peptide for 48 hours, control library-transduced cells exhibited pronounced cell spreading and membrane ruffling indicative of actin remodeling, whereas *G*_α_*q*-knockout cells failed to respond. **(B)** Representative FACS plots of mixed populations containing *G*_α_*q*-knockout (BFP^+^) and CRISPR library control (GFP^+^) cells treated with AKH peptide for 0, 24, 48, and 72 hours. A progressive decrease in GFP^+^ cells and enrichment of BFP^+^/DAPI^+^ cells over time indicates selective depletion of signaling-competent cells and survival of G_α_q-deficient cells under prolonged AKH stimulation. **(C)** Schematic of the pooled CRISPR-Cas9 screen. AkhR-expressing S2R+ cells were mixed in a 1:1 ratio: BFP-labeled *G*_α_*q* knockout cells and GFP-labeled genome-wide CRISPR library-transduced cells. Populations were either left untreated or exposed to AKH peptide for selection. Screens were independently conducted in parallel. **(D)** Genes identified in the screen were ranked using the robust rank aggregation (RRA) score calculated by MAGeCK analysis. Top-ranking candidate genes are color-coded by functional category**. (E)** Clustered interaction matrix of AlphaFold-Multimer–predicted protein-protein interactions among top 50 CRISPR screen hits. The matrix shows pairwise interaction scores for high-confidence structural predictions. Color scale indicates predicted interface lDDT (iLDDT) scores, and dot size reflects average confidence (iLDDT_avg) across interaction models. Hub proteins (e.g., ZDHHC8, Su(var)3–3) form dense interaction clusters, suggesting functional modules within the AkhR signaling network. **(F)** Bar graph summarizing the top 15 proteins with the highest number of high-confidence PPIs from AFM predictions, highlighting potential signaling hubs such as ZDHHC8, Su(var)3–3, and Hipk. **(G)** Network diagram of selected high-confidence PPIs predicted by AFM. Red nodes represent central hub proteins with multiple predicted interactions; edges indicate inferred structural associations. ZDHHC8 and CG5447 (Golgin A7) emerge as tightly connected components in the AkhR signaling network.

Analysis using MAGeCK identified several categories of genes whose loss promoted conferred a selective enrichment under AKH treatment **(Figure 2D).** These included (1) Core Gαq pathway components such as *AkhR*, *G*β*13F*, *G*γ*1*, *Gaq*, *CG7650* (fly homolog of phosducin) and *rush hour (rush)* which interacts with GPCR-mediated phosphatidylinositol 3-phosphate (PI3P), validating the sensitivity of the screen (2) MAPK/ERK and Rho pathways e.g., *Mekk1, licorne (lic), Rho Kinase (Rok), and tribble (trbl)*, suggesting cytoskeletal and stress-response crosstalk (3) Epigenetic modifiers: including *[histone H3]-lysine(36) Nuclear receptor binding SET domain protein* (*Mes-4)*, *CG8026*, and *CG4747*, potentially linking sustained GPCR signaling to transcriptional adaptation, and (4) Pro-apoptotic factors: such as ER membrane protein complex subunit 6 (*CG11781)*, indicating a signaling-to-death axis downstream of AkhR. In addition, the screen identified two uncharacterized genes *CG34449* and *CG5447*. *CG34449* is predicted to encode fly homolog of human ZdhhC8^16^, a palmitoyl transferase, and CG5447 is predicted to encode the fly homolog of human Golga7A, previously reported as a ZdhhC cofactor^13^.

### AlphaFold-Multimer Structural Predictions Reveal Functional Clusters

To explore potential physical interactions among the top candidates from our genome-wide CRISPR screen, we applied AlphaFold-Multimer (AFM) to predict protein–protein interactions (PPIs) across the top 50 genes ranked by MAGeCK. This cutoff was chosen to enrich for likely functional hits while maintaining computational feasibility and interpretability. We performed an all-by-all pairwise prediction (50 × 50), excluding large multimeric complexes incompatible with current modeling constraints.

Predicted interaction scores are summarized as a heatmap **(Figure. 2D)**, in which blue-shaded cells denote high-confidence PPIs^17^. In parallel, a frequency bar plot **(Figure. 2E)** quantifies how often individual proteins appeared in predicted complexes, highlighting interaction hubs such as Su(var)3–3, ZDHHC8, and Hipk. To visualize global interaction architecture, we projected the pairwise predictions into a low-dimensional space. This analysis revealed that top hits such as ZDHHC8, CG5447, and Gβ13F resolve as a distinct cluster, suggesting specificity and tight functional coupling. By contrast, many remaining genes appear in a more densely packed region, limiting individual resolution.

The resulting predictions revealed numerous high-confidence interactions within and across defined functional categories, including Gαq signaling components, palmitoylation machinery, MAPK/Rho kinases, cytoskeletal regulators, and chromatin modifiers. For instance, AFM confidently predicted canonical complexes such as Gαq–Gβ13F^18^ and Gβ13F–Gγ1, known components of the heterotrimeric G protein complex. Importantly, the model also recovered a direct interaction between AkhR and Gαq, consistent with their known receptor–effector relationship and validating the ability of AFM to capture biologically relevant GPCR–G protein complexes^19^. Similarly, ZDHHC8 and CG5447 (Golgin A7), which form a known palmitoyltransferase–adaptor pair, were also predicted to interact with high structural confidence, further substantiating the analysis. These expected interactions serve as internal positive controls, supporting the reliability of novel predictions derived from the same dataset.

In addition to intra-category connections, AFM also predicted cross-functional associations, for example, between MAPK signaling kinases and chromatin remodelers, or between apoptotic proteins and palmitoylation enzymes. These results suggest possible scaffolding or feedback mechanisms linking membrane signaling to transcriptional and cytoskeletal programs during sustained AkhR–Gαq activation.

Together, the integration of AFM-based structure prediction with our functional CRISPR screen uncovers both expected and novel regulators of Gαq-mediated signaling^17^. This dual-layer approach provides a structural framework for interpreting functional clusters and identifying candidate protein complexes mediating glucagon-like GPCR activity, laying a foundation for future mechanistic dissection.

### ZDHHC8 and Golgin A7 Promote AkhR–G**α**q Signaling via Palmitoylation-Dependent Stabilization

Among the AFM results, ZDHHC8 (CG34449) emerged as one of the top proteins forming numerous high-confidence protein-protein interactions (PPIs). Notably, its known functional partner, Golgin A7 (CG5447), also ranked highly in PPI connectivity, reinforcing the significance of the palmitoylation machinery in AkhR signaling. Based on these findings, we prioritized ZDHHC8 and Golgin A7 for functional validation. Independent knockouts of *CG34449* and *CG5447* in AkhR-expressing S2R^+^ cells conferred strong resistance to AKH-induced enrichment, phenocopying the effect of *AkhR* knockout **(Figure 3A).** These cells remained failed to undergo actin collapse or the morphological remodeling characteristic of signaling-competent cells, supporting a key role for protein palmitoylation in sustaining Gαq-mediated signaling and actin remodeling-mediated enrichment responses.

**Figure 3.**
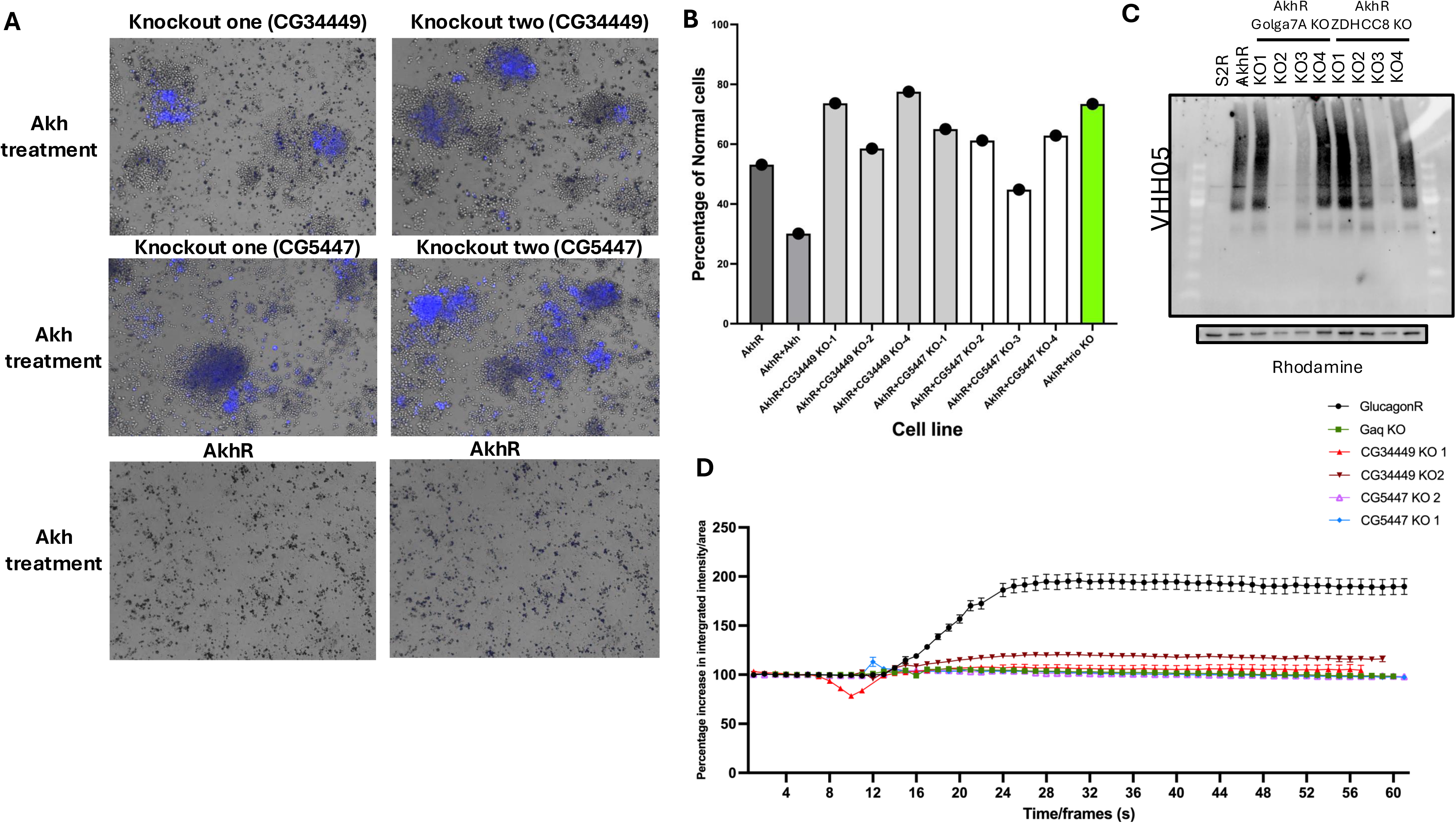
ZDHHC8 (CG34449) and Golgin A7 (CG5447) are required for AkhR-mediated G_α_q signaling, receptor stability, and cellular responses. **(A)** Representative brightfield images of S2R^+^ cells expressing AkhR following AKH stimulation show widespread cytopathic effects in wild-type cells, which are suppressed in *CG34449* and *CG54*47 knockout lines. Blue overlay represents BFP-expressing sgRNA transfected cells **(B)** Quantification of calcium mobilization responses following AKH stimulation in control and knockout lines using Fluo-4 AM imaging. Both *CG34449* and *CG5447* knockouts exhibit a significant reduction in cytosolic calcium elevation compared to AkhR-expressing control cells, comparable to G_α_q loss-of-function **(C)** Immunoblot analysis of AkhR protein levels in AkhR-expressing cells upon CRISPR knockout of CG34449, CG5447 in different clones. Loss of CG34449 or CG5447 causes a marked reduction in AkhR abundance in few clones. Actin is shown as loading control **(D)** Whole-body triacylglycerol (TAG) levels in adult flies with fat body–specific *CG5447* knockdown are significantly elevated compared to wild-type (*w*^1118^) and mirror the metabolic phenotype of *AkhR* knockdown **(E)** Time-course imaging of Fluo-4 calcium flux in response to AKH stimulation in indicated genotypes. Points represent mean fluorescence intensity across cells, showing blunted calcium signaling in *CG34449* and *CG5447* knockouts

Quantification of cell complexity via SSC/FSC flow cytometry confirmed reduced structural remodeling in *CG34449* and *CG5447* knockout cells compared to controls **(Figure 3B),** consistent with impaired receptor signaling. Further, Western blot analysis revealed that RNAi-mediated depletion of either *CG34449* or *CG5447* caused a marked reduction in AkhR protein levels **(Figure 3C),** similar to *AkhR* knockdown (**Supplementary Figure 1B**), suggesting that these genes are required for maintaining AkhR protein levels. In line with these observations, calcium imaging using Fluo-4 AM demonstrated that AKH-induced Ca²^+^ mobilization was significantly blunted in both *CG34449* and *CG5447* knockouts, comparable to the effect seen in Gαq-deficient cells **(Figure 3D)**, supporting a critical role for these genes in AkhR– Gαq pathway activation.

### ZDHHC8-Dependent Palmitoylation of G**α**q Revealed by Click Chemistry Labeling

We next investigated the mechanism by which ZDHHC8 regulates the AkhR/Gαq pathway. DHHC-type palmitoyl acyltransferases (PATs) are zinc-finger membrane enzymes defined by a conserved Asp-His-His-Cys catalytic motif, which attach the 16-carbon fatty acid palmitate to specific cysteine residues on target proteins. This family comprises 23 enzymes in humans and mediates reversible S-palmitoylation, dynamically regulating the localization and function of numerous signaling proteins analogously to phosphorylation^20–22^. Given that ZDHHC8 is an enzyme that adds palmitate lipids to cysteine residues on target proteins, we hypothesized that a key substrate of ZDHHC8 in this pathway might be either the receptor itself or the Gαq protein. Many GPCRs, especially Family A receptors, have a cysteine in their C-terminal tail that can be palmitoylated, which affects receptor localization and signaling. The AkhR sequence contains two cysteine residues in its C-terminus that could potentially be palmitoylation sites. On the other hand, mammalian Gαq is known to be palmitoylated on its N-terminal cysteine as a mechanism for membrane attachment^23^. To distinguish which substrate is functionally relevant for ZDHHC8, we assessed palmitoylation of Gαq and AkhR in the presence or absence of ZDHHC8.

We first directly measured Gαq palmitoylation using metabolic labeling with the palmitate analog 17-octadecynoic acid (17-ODYA) followed by click chemistry detection with TAMRA-azide^24^ **(Figure 4A)**. S2R+ cells transiently expressing 3X-FLAG-tagged Gαq were incubated with 17-ODYA, and after 24 hours, Gαq was immunoprecipitated and subjected to copper-catalyzed azide–alkyne cycloaddition (CuAAC) to attach the fluorescent TAMRA-azide tag. SDS-PAGE and in-gel fluorescence analysis revealed robust labeling of Gαq in wild-type cells, confirming that Gαq is palmitoylated under basal conditions **(Figures 4B)**. Strikingly, in *Zdhhc8* knockout cells, the TAMRA fluorescence signal for Gαq was almost completely lost, despite equal expression of total Gαq protein. Quantification indicated that ZDHHC8 loss reduced Gαq palmitoylation by ∼70–80%, demonstrating that ZDHHC8 is the predominant enzyme responsible for Gαq palmitoylation.

**Figure 4.**
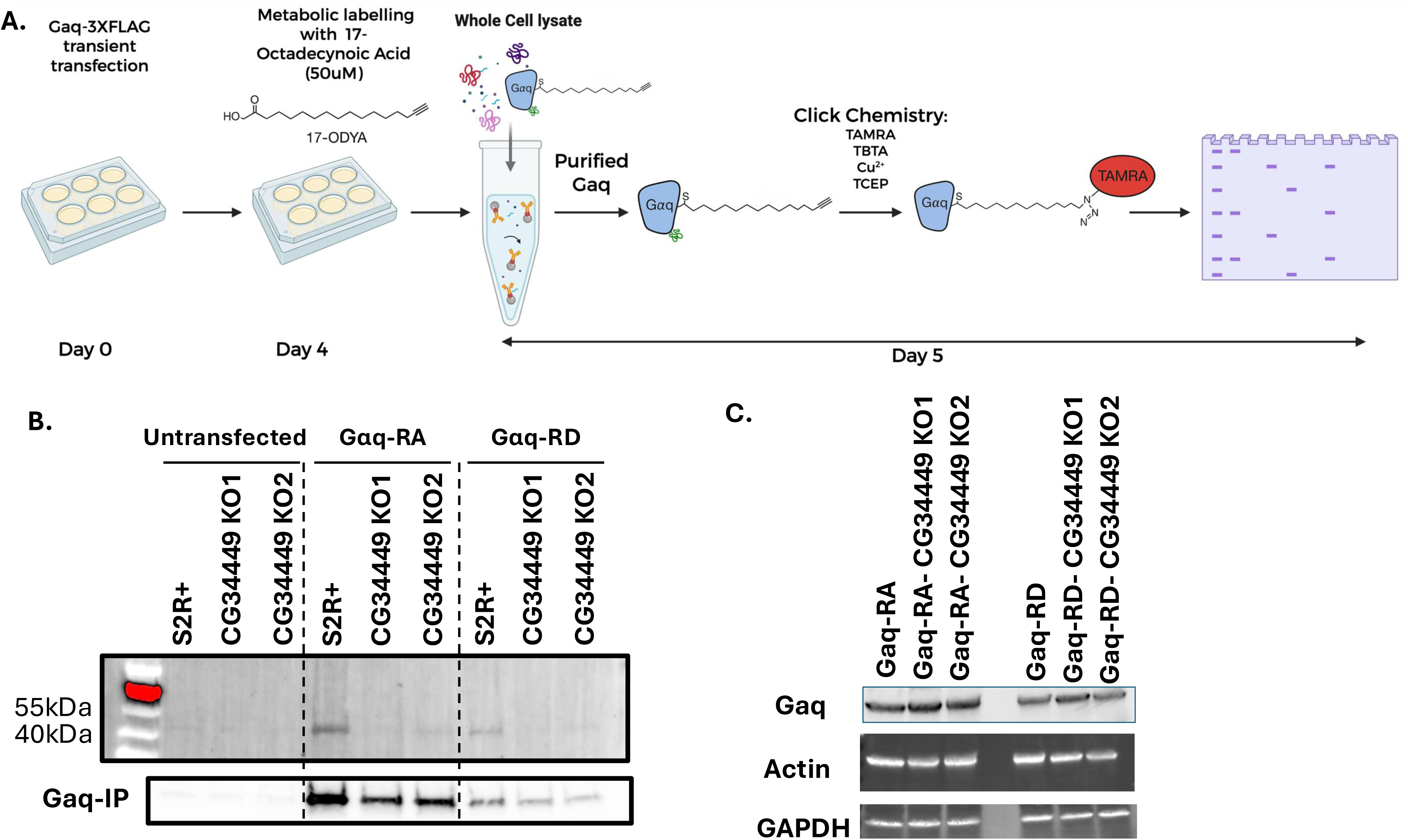
G_α_q is palmitoylated in S2R^+^ cells via a ZDHHC8-dependent mechanism. **(A)** Schematic of the experimental workflow for detecting G_α_q palmitoylation using copper-catalyzed azide–alkyne cycloaddition (CuAAC) click chemistry. S2R^+^ cells were transiently transfected with *G*_α_*q-3xFLAG* and metabolically labeled with 17-octadecynoic acid (17-ODYA, 50 _μ_M) for 24 h. FLAG-tagged G_α_q was immunoprecipitated and subjected to click chemistry with TAMRA-azide in the presence of TBTA, Cu²^+^, and TCEP. Samples were resolved by SDS-PAGE and analyzed by in-gel fluorescence and western blotting. **(B)** G_α_q palmitoylation is markedly reduced in *CG34449* (*Zdhhc8*) knockout S2R^+^ cells compared to wild-type, indicating that ZDHHC8 is required for G_α_q palmitoylation. Two independent KO clones (KO1 and KO2) show consistent loss of TAMRA signal. Lower panels show total FLAG-G_α_q input as loading control. **(C)** Western blot analysis of total G_α_q expression in wild-type and CG34449 (ZDHHC8) knockout S2R^+^ cells. Cells were transiently transfected with either G_α_q-RA or G_α_q-RD splice isoforms and analyzed in two independent ZDHHC8 knockout clones (KO1 and KO2). No significant change in G_α_q protein levels was observed upon ZDHHC8 loss. Actin and GAPDH serve as loading controls.

Importantly, total Gαq protein levels remained unchanged in ZDHHC8 knockout cells across two independent Gaq isoforms (**Figure 3C**), demonstrating that the loss of palmitoylation does not alter Gαq protein abundance. This finding suggests that ZDHHC8-dependent palmitoylation regulates Gαq function and localization rather than its stability. This observation is consistent with prior studies in mammalian systems, where palmitoylation was shown to control protein trafficking or activity without necessarily affecting protein levels^25, 26^. These results collectively establish ZDHHC8 as the principal palmitoyltransferase responsible for Gαq lipidation and support a model in which palmitoylation governs Gαq membrane targeting and signal competence without altering its expression.

### ZDHHC8 and Golgin A7 are required for the membrane localization of AkhR

As mammalian GPCRs have been shown to be palmitoylated, we next asked whether ZDHHC8 or Golgin A7 might additionally act on AkhR. Immunofluorescence imaging revealed that in control cells, AkhR localized primarily to the plasma membrane. However, *CG34449* knockout led to intracellular accumulation of AkhR **(Figure 5A)**. Co-localization studies showed that internalized AkhR failed to overlap with Golgi (Golgin245), ER (Calnexin 99A), or early endosome (Hrs) markers, but co-localized with lysosomes (LysoTracker) **(Figure 5B)**, suggesting that the internalized receptor may be degraded in the lysosome.

**Figure 5.**
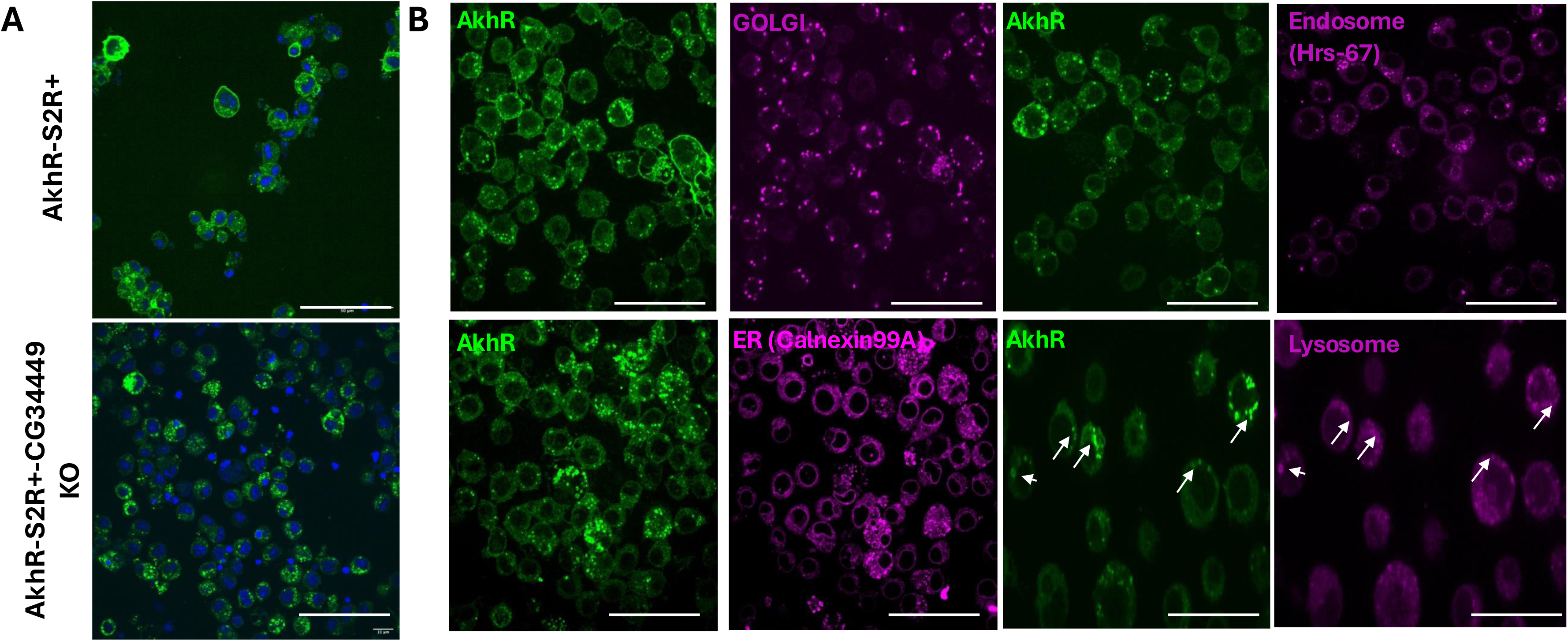
ZDHHC8 is essential for proper plasma membrane localization of AkhR; its loss results in receptor misrouting to lysosomes. **(A)** Immunofluorescence images of S2R^+^ cells stably expressing AkhR (green) in either wild-type background (top) or *Zdhhc8* knockout (*CG34449* KO; bottom). In wild-type cells, AkhR shows prominent localization at the plasma membrane. In contrast, *CG34449* KO cells display marked intracellular accumulation of AkhR, consistent with trafficking defects. Nuclei are stained with DAPI (blue). **(B)** Co-localization analysis of AkhR (green) with organelle-specific markers (magenta) in S2R^+^ cells. In *CG34449* KO background, AkhR fails to co-localize with Golgi (Golgin245), ER (Calnexin99A), or early endosomes (Hrs), but shows strong overlap with LysoTracker-labeled lysosomes (arrows), indicating redirection of mislocalized AkhR to lysosomal degradation. Each pair of panels shows the same field imaged for AkhR and the respective organelle marker. Scale bars???

Mammalian GPCRs have been shown to be palmitoylated on Cysteine residues at the C-terminus [1,2]. Sequence alignment predict that these sites are conserved in *Drosophila* AkhR (C426 and C434) (**Figure 6A)**. To test their importance, we generated cysteine-to-alanine single (C426A, C434A) and double (C426A/C434A) point mutants and stably expressed them in S2R^+^ cells. Upon AKH stimulation, cells expressing wild-type AkhR exhibited dynamic membrane ruffling and spreading. By contrast, all three cysteine mutants failed to respond morphologically to ligand stimulation, suggesting a loss of receptor activity **(Figure 6B)**.

**Figure 6.**
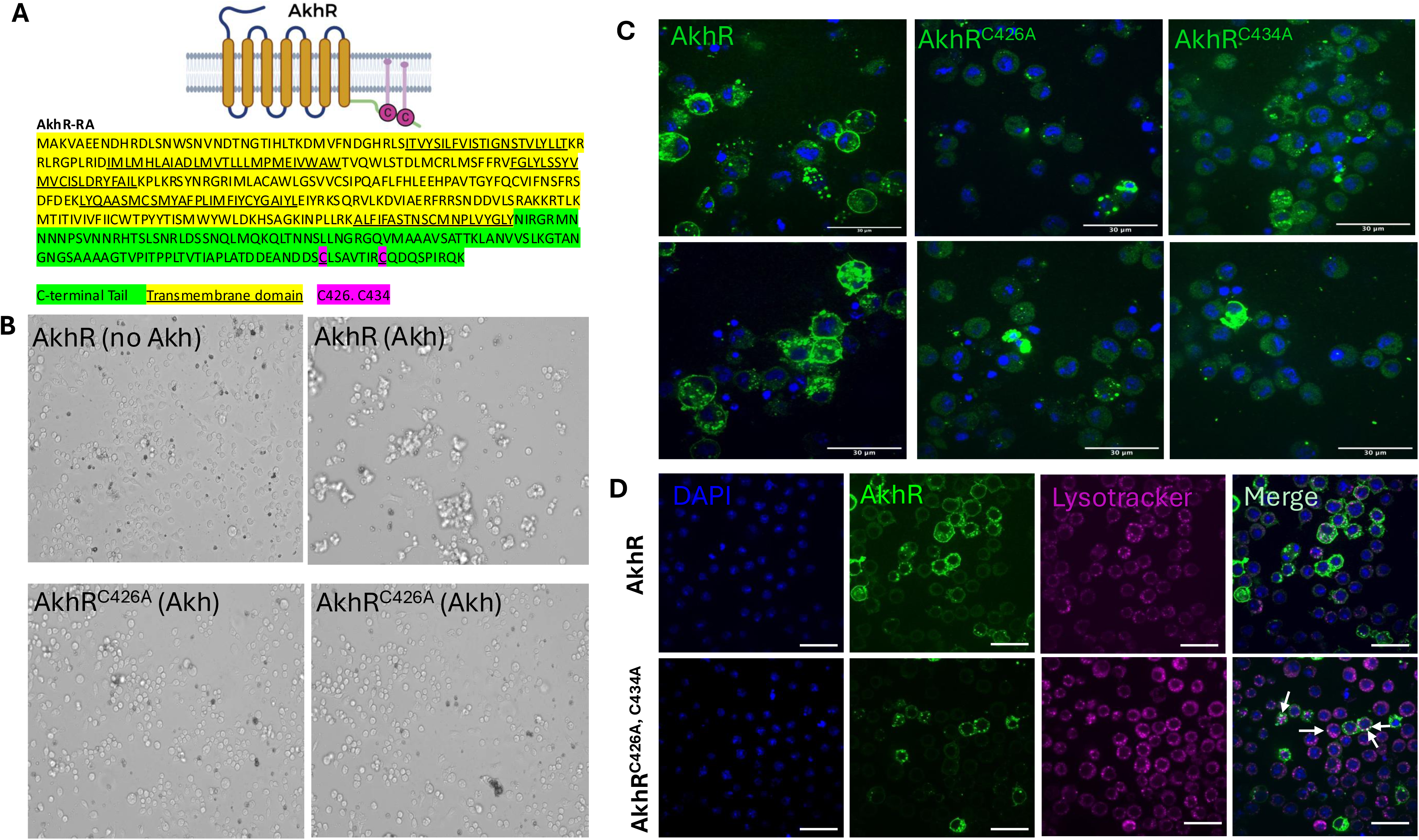
Conserved C-terminal cysteine residues regulate AkhR surface stability and lysosomal degradation. **(A)** Amino acid sequence of the AkhR isoform used in this study, highlighting predicted transmembrane domains (yellow), C-terminal peptide (green), and C-terminal cysteine residues C426 and C434 (pink), which are candidate palmitoylation sites. **(B)** Brightfield imaging of S2R^+^ cells expressing wild-type AkhR or cysteine-to-alanine point mutants (C426A and C434A) following AKH peptide treatment. While wild-type AkhR-expressing cells display characteristic membrane ruffling in response to ligand stimulation, all mutant variants fail to undergo morphological changes, indicating a loss of signaling competence. **(C)** Immunofluorescence staining reveals markedly reduced AkhR protein levels in cells expressing C426A and C434A mutants compared to wild-type AkhR. Mutant-expressing cells show punctate intracellular staining, suggesting receptor destabilization or mislocalization. **(D)** Co-localization of AkhR and LysoTracker in S2R^+^ cells expressing the C426A/C434A double mutant (indicated by white arrows). Unlike wild-type AkhR, which localizes predominantly to the plasma membrane, the double mutant is retained in intracellular compartments and overlaps extensively with lysosomal markers, indicating lysosome-mediated degradation. Scale bars: 30 µm.

We next assessed whether these mutations affect AkhR protein levels or trafficking. Immunofluorescence revealed that the point mutants exhibited substantially lower receptor levels than wild-type AkhR, with the C426A/C434A double mutant showing the most dramatic reduction **(Figure 6C)**. Notably, while wild-type AkhR localized predominantly to the plasma membrane, the double mutant accumulated intracellularly in punctate compartments. Co-staining with LysoTracker revealed strong colocalization of the double mutant with lysosomes **(Figure 6D)**, suggesting that in the absence of these cysteine residues, AkhR may be degraded in the lysosome.

These observations phenocopy the effects of ZDHHC8 or Golgin A7 loss and suggest that C426 and C434 serve as palmitoylation sites essential for maintaining AkhR surface stability. The requirement for these residues is consistent with the known role of palmitoylation in promoting plasma membrane localization, receptor stability, and resistance to lysosomal degradation [1,3].

**Figure 7.**
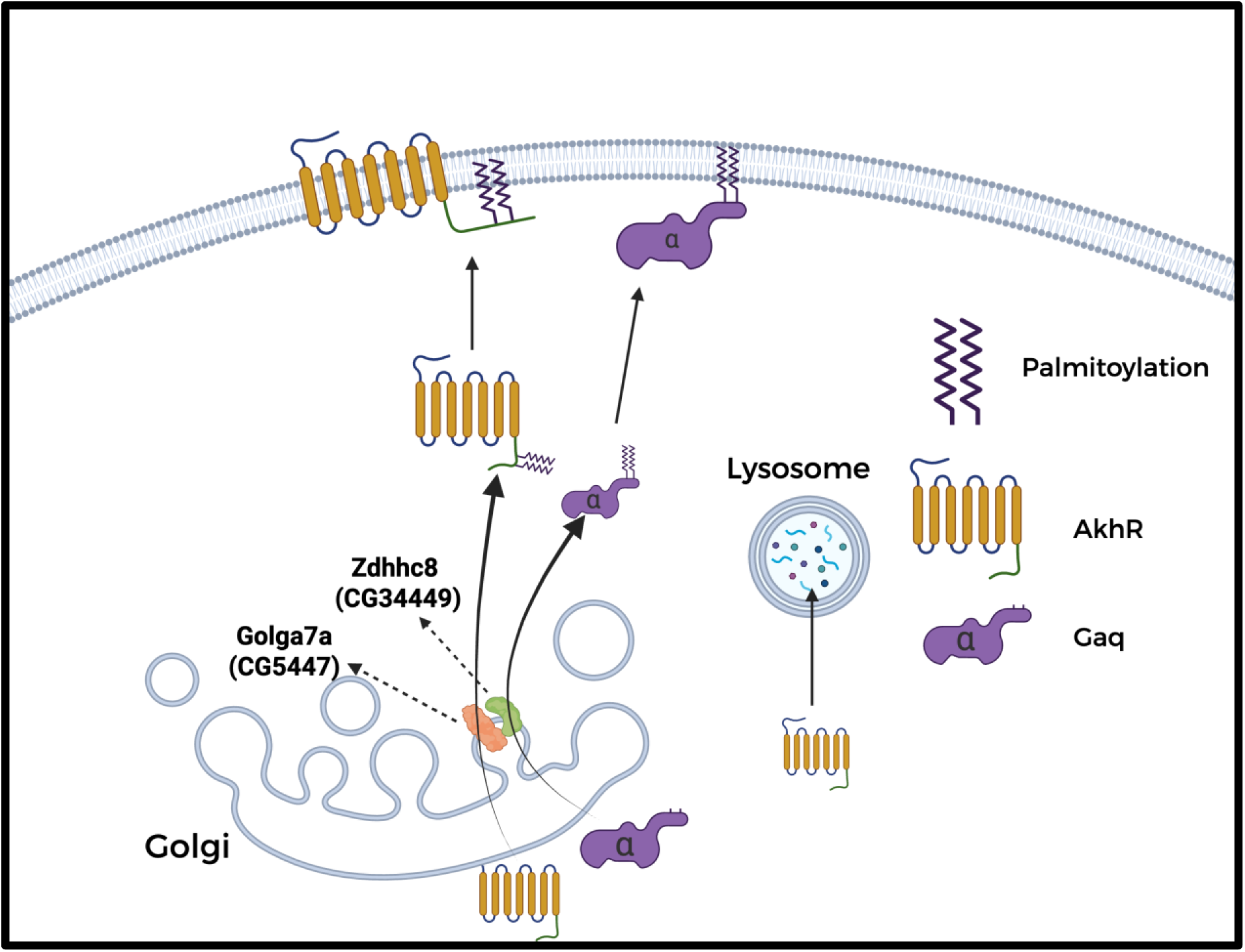
Model of ZDHHC8-dependent palmitoylation and regulation of the AKH (dGlucagon)–G_α_q signaling axis in *Drosophila*. Schema illustrating the role of ZDHHC8 (CG34449) and its cofactor Golga7a (CG5447) in regulating the membrane localization and signaling competence of G_α_q and the AKH receptor (AkhR). G_α_q and AkhR are synthesized and trafficked through the Golgi apparatus. The palmitoyl acyltransferase ZDHHC8 (CG34449) and its adaptor Golga7a localize at the Golgi and mediate S-palmitoylation of G_α_q, a modification required for its membrane attachment. Palmitoylated G_α_q is then able to associate with AkhR at the plasma membrane to enable efficient GPCR signaling. In the absence of ZDHHC8 or Golga7a, G_α_q fails to undergo palmitoylation, leading to reduced membrane localization and impaired coupling to AkhR. Concurrently, AkhR itself most likely requires palmitoylation of conserved C-terminal cysteine residues for proper plasma membrane localization. Loss of *Zdhhc8*, *Golga7a*, or the receptor’s palmitoylation sites leads to mislocalization of AkhR and its subsequent degradation via the lysosomal pathway, ultimately attenuating signaling output. These results define a dual palmitoylation-dependent mechanism that governs both receptor and G protein stability and function in a metabolic GPCR pathway.

## Discussion

This study identifies ZDHHC8 as a central palmitoyltransferase required for the activation of the Gαq-coupled adipokinetic hormone receptor (AkhR) signaling axis in *Drosophila*. Palmitoylation of mammalian Gα subunits, including Gαs, Gαi, and Gαq, has been previously reported and shown to be important for membrane association and signaling activity. However, the specific palmitoyltransferases responsible, and whether a single DHHC enzyme can simultaneously regulate both a GPCR and its downstream Gα subunit, have not been previously demonstrated. Our study identifies ZDHHC8 as the principal enzyme mediating Gαq palmitoylation in *Drosophila* and reveals a dual-substrate mechanism that may be conserved across GPCR signaling modules. We show that CG34449 (ZDHHC8) is essential for the palmitoylation and function of both the GPCR AkhR and its downstream G protein Gαq, two core components of a canonical Class A GPCR signaling module. While previous studies have demonstrated that either G proteins or GPCRs can be palmitoylated by DHHC-family enzymes^22, 26, 27^, this study presents, to our knowledge, the first demonstration that a single ZDHHC enzyme putatively governs the palmitoylation of both the receptor and its coupled G protein within the same pathway. This dual regulation adds a new layer of mechanistic insight into how GPCR signal propagation and fidelity may be coordinated at the level of lipid post-translational modification.

Rather than providing fine-tuned control over signaling, the convergence of both substrates on a single enzyme likely enforces a binary checkpoint i.e. ZDHHC8 activity may function as a molecular gate that either permits or blocks signal transduction by regulating the membrane localization of both AkhR and Gαq. This mechanism may minimize differential modulation between the receptor and G protein, potentially enhancing the robustness and coordination of signaling outputs.

Our data indicate that AkhR may require palmitoylation for both stability and surface localization. We attempted to directly detect AkhR palmitoylation using immunoprecipitation–mass spectrometry (IP-MS) and click chemistry. However, these approaches were unsuccessful due to experimental limitations: the receptor is heavily glycosylated and contains few lysine and arginine residues, which limited proteolytic digestion and peptide coverage in MS analysis. In the case of click chemistry, glycosylation likely impaired efficient labeling or immunoprecipitation, and the resulting high molecular weight smear on western blots prevented reliable overlap of the TAMRA signal with the AkhR band, further complicating interpretation. Due to these experimental limitations, our evidence for AkhR palmitoylation is currently indirect, based on mutagenesis of conserved cysteine residues and lysosomal degradation phenotypes. While these findings suggest that C-terminal palmitoylation governs AkhR localization fate, definitive biochemical confirmation (e.g., acyl-exchange assays or mass spectrometry) will be needed to validate AkhR as a bona fide palmitoylation substrate. Conversely, our 17-ODYA metabolic labeling and click chemistry experiments provide direct evidence that Gαq is palmitoylated in a ZDHHC8-dependent manner, supporting a model where ZDHHC8 is the principal enzyme controlling Gαq lipidation and plasma membrane localization.

This dual-substrate control may have broader implications beyond *Drosophila*. Many Class A GPCRs in mammals, including the β2-adrenergic receptor, rhodopsin, and the glucagon receptor (GCGR), are palmitoylated on their cytoplasmic tails, a modification known to influence receptor stability, trafficking, and coupling. Similarly, mammalian Gαq is palmitoylated at its N-terminal cysteine, which is required for membrane anchoring and efficient effector interaction. While enzymes such as ZDHHC5, ZDHHC7, and ZDHHC9 have been implicated in GPCR or G protein palmitoylation individually, the coordinated control of both components by a single DHHC enzyme has not been previously reported. Our study thus raises the possibility that a conserved subset of DHHCs (e.g., ZDHHC8) may act as master regulators of entire GPCR signaling modules, a hypothesis that warrants testing in mammalian systems.

ZDHHC8 itself has drawn increasing attention in human disease. It is highly expressed in the brain, genetically linked to schizophrenia risk, and implicated in AMPA receptor trafficking via palmitoylation of neuronal scaffolds. Moreover, emerging studies suggest roles for palmitoylation in oncogenic GPCR signaling and metabolic regulation. As such, the ZDHHC8–Golgin A7 complex may represent a convergence point for modulating GPCR activity in diverse physiological and pathological contexts. Our results support the idea that targeting palmitoylation enzymes, either pharmacologically or genetically, could provide a lever to rewire or dampen aberrant GPCR signaling. While general palmitoylation inhibitors like 2-bromopalmitate (2-BP) lack specificity, recent advances in designing DHHC-specific inhibitors or substrate-competitive peptides raise the prospect of selectively targeting individual PATs like ZDHHC8.

Future work should test whether the ZDHHC8–Gαq–GPCR regulatory axis observed here is conserved in mammalian glucagon receptor signaling, and whether tissue-specific ZDHHC8 activity correlates with physiological demands for Gαq-coupled hormone responses. It will also be critical to determine how dynamic palmitoylation of GPCR components interacts with other modes of regulation, such as phosphorylation, ubiquitination, and endocytosis, and whether such crosstalk defines context-specific outputs in metabolic or neurological tissues.

In conclusion, our study defines a previously uncharacterized regulatory axis in GPCR signaling, wherein ZDHHC8-dependent palmitoylation coordinates the membrane localization and stability of both a receptor and its G protein. This mechanism enforces signaling competence through lipid modification and establishes a model for how single enzymes can regulate multicomponent signaling cascades. By integrating genetic, biochemical, and imaging approaches, we uncover a dual palmitoylation system that may be widely conserved and therapeutically actionable in GPCR-associated diseases.

## Acknowledgements

We thank Richard Binari for general lab assistance. BioRender was used to make a subset of figure panels. NP was supported by NIH grants R01GM084947, R01GM067761, R24OD019847. MA was supported by the American Heart Association (Award ID 24PRE1189954). NP is an investigator of the Howard Hughes Medical Institute. Portions of the manuscript’s text were drafted with the assistance of ChatGPT (OpenAI), under the supervision of the authors. All content was reviewed and edited by the authors for accuracy and originality This article is subject to HHMI’s Open Access to Publications policy. HHMI lab heads have previously granted a nonexclusive CC BY 4.0 license to the public and a sublicensable license to HHMI in their research articles. Pursuant to those licenses, the author-accepted manuscript of this article can be made freely available under a CC BY 4.0 license immediately upon publication.

## Materials and Methods

### Cell Lines and Culture Conditions

*Drosophila* S2R^+^ cells (obtained from the *Drosophila* RNAi Screening Center, DRSC) were used for all experiments. Cells were maintained at 25°C in Schneider’s *Drosophila* medium (Thermo Fisher Scientific) supplemented with 10% heat-inactivated fetal bovine serum (FBS) and 1× penicillin-streptomycin. Cultures were grown in T-75 flasks without COD. For routine subculture, cells were passaged at a 1:5 ratio every 3–4 days to maintain logarithmic growth. S2R^+^ derivative line “PT5” (NPT005) was used as the parental cell line for genome engineering. This line contains an attP site for ΦC31 integrase-mediated transgene insertion (facilitating stable library integration, see below) and was further engineered to stably express Cas9 (see below). All cell lines were routinely tested for mycoplasma contamination and confirmed negative.

### Stable Cell Line Generation (Cas9 and GPCR Expression)

To enable genome-wide CRISPR screening, we generated a stable S2R^+^-Cas9 cell line. S2R^+^ PT5 cells were transfected with a plasmid encoding a *Drosophila*-codon-optimized Cas9 under a metallothionein promoter (pMK33/Cas9). Selection was performed in 200 µg/mL Hygromycin B for 4–8 weeks, yielding a polyclonal line stably expressing Cas9. The resulting S2R^+^-Cas9 cells were maintained in Schneider’s medium + 10% FBS with 200 µg/mL Hygromycin to ensure continued Cas9 expression until introduction of the sgRNA library.

For functional assays, we established stable S2R^+^ cell lines overexpressing the Adipokinetic hormone receptor (AkhR) or the Capa receptor (CapaR). Coding sequences for *Drosophila AkhR* (*CG11325*) and *CapaR* (*CG14575*) were cloned into a pAc5-STABLE expression vector (derived from pAc5.1) with an N-terminal VHH05 epitope tag to facilitate detection. The VHH05 tag is a camelid single-domain antibody epitope that was recognized by a specific monoclonal antibody in immunoassays^28^. S2R^+^-Cas9 cells were transfected with the *AkhR-VHH05* or *CapaR-VHH05* plasmid along with a neomycin-resistance marker. After 48 hours, selection was applied using 1 mg/mL G418 (Thermo Fisher) for ∼3 weeks. Surviving cells were dilution-cloned in 96-well plates to isolate single-cell colonies. Clonal lines with robust receptor expression were identified by Western blotting with anti-VHH05 antibody and maintained as stable AkhR-expressing and CapaR-expressing S2R^+^. All experiments involving AkhR or CapaR stimulation were performed using these clonal stable cell lines.

### Genome-wide pooled CRISPR/Cas9 Screen

We performed a pooled genome-wide CRISPR knockout screen in the AkhR-expressing S2R^+^-Cas9 cells to identify genes required for AkhR/Gαq-mediated signaling. The *Drosophila* genome-wide sgRNA library was the version 2 library described in previous study^29^. Briefly, the library contains ∼85,000 sgRNAs (6 guides per gene) targeting ∼14,000 *Drosophila* genes, cloned in the pLib6.4 plasmid (which carries *Act5C-GFP-T2A-Puro* for selection). For library delivery, we utilized site-specific ΦC31 integrase-mediated integration into the attP site of the S2R^+^ PT5 cells along with AcrIIa4 (IntAC) to delay editing events until the completion integration^29^. AkhR-expressing S2R^+^-Cas9 cells were co-transfected with the pLib6.4 sgRNA library plasmid DNA and IntAC plasmid at a 1:1 ratio using Effectene transfection reagent (Qiagen), following the manufacturer’s protocol. Cells were plated such that at least 1,500 cells were transfected per sgRNA to ensure adequate library coverage. Two days post-transfection, selection for library integrants was applied by adding puromycin (5 µg/mL) to the culture medium. Cells were grown under puromycin selection for 5–7 days, during which the pooled population was expanded to maintain representation of all sgRNAs (maintaining >1500 cells per sgRNA at each passage).

For the functional selection, we exploited the fact that chronic activation of AkhR triggers cytotoxic Gαq signaling in wild-type cells, whereas cells lacking essential signaling components survive. We therefore performed a competitive co-culture screen. In parallel with the library-transduced population, we generated a reference population of “signal-deficient” cells by knocking out the gene encoding Gαq (encoding the G protein αq subunit required for AkhR signaling) in the *AkhR* cell line. This was achieved by transfecting a plasmid expressing an sgRNA targeting *galphao* (*G*α*q*) along with a blue fluorescent protein marker (BFP). Single-cell cloning yielded a *G*α*q*-knockout clonal line, which was labeled with BFP (via a constitutive expression cassette). We mixed the GFP^+^ library-transduced cells and BFP^+^ *G*α*q*-knockout cells at a 1:1 ratio (∼5×10^6 cells each) to create the screening population. Two parallel populations were set up i.e. one was maintained in normal medium (unselected control), and the other was subjected to AkhR agonist selection. For selection, synthetic *Drosophila* Adipokinetic Hormone peptide (AKH; sequence QLTFSPDW-amide) was added to the culture at 10 µg/mL. Fresh AKH was replenished every 48 hours for a prolonged stimulation period of 8 days. Throughout this period, cells were maintained in log-phase growth by diluting or splitting as needed (with AKH continuously present in the selected population). The AKH concentration and duration were chosen based on preliminary dose-response tests to induce significant **enrichment** effects in wild-type AkhR cells without eliminating the culture.

During the selection, the relative abundance of GFP^+^ (library) versus BFP^+^ (*G*α*q*-null) cells was monitored every 2 days by flow cytometry (see Flow Cytometry below). As expected, chronic AKH stimulation led to a progressive loss of GFP^+^ library cells and an enrichment of BFP^+^ *G*α*q*-deficient cells, indicating selective depletion of signaling-competent (AKH-sensitive) cells. After 8 days of AKH treatment, the enriched cell populations (AKH-treated and control) were harvested for genomic DNA extraction using a Qiagen Blood & Cell Culture DNA kit. sgRNA sequences were PCR-amplified from genomic DNA in two steps (to add flow-cell adapters and indices) as previously described. The resulting amplicon libraries were deep sequenced on an Illumina platform to determine sgRNA representation in each condition.

#### Screen Data Analysis

Sequencing reads were mapped to the sgRNA library index. Read counts per sgRNA in AKH-treated vs. control samples were normalized and analyzed using MAGeCK (Model-based Analysis of Genome-wide CRISPR-Cas9 Knockout) software^30^. MAGeCK was run in robust rank aggregation mode to identify genes enriched or depleted under AKH selection. In this analysis, sgRNAs that confer an enrichment advantage under AKH (i.e., target genes required for AKH-induced cytotoxic signaling) show increased abundance in the AKH-treated sample relative to control. MAGeCK assigns each gene an enrichment score and false discovery rate (FDR) based on the consistency of sgRNA rank changes. Genes were ranked by MAGeCK’s robust rank aggregation (RRA) scorefile, and those with FDR < 0.1 were considered high-confidence hits for regulating AkhR/Gαq signaling (**Supplementary File 1**). Downstream pathway analysis and functional categorization of top hits were conducted using FlyBase and Gene Ontology databases.

### AlphaFold-Multimer Predictions

To predict protein–protein interactions among genes identified in the genome-wide CRISPR screen, we employed LocalColabFold v.1.5.5^31^, which integrates AlphaFold-Multimer (AFM) v.2.3.1^31^ and utilizes MMseqs2 (14-7e284 version) for generating multiple sequence alignments. The input set consisted of the top 50 genes ranked by MAGeCK analysis (FDR < 0.1), which were selected to balance enrichment strength and computational feasibility. Their longest isoforms were retrieved from FlyBase and used for predictions.

We performed an exhaustive pairwise all-by-all screen, generating 1,225 unique heterodimeric combinations (**Supplementary File 2**). We predicted five AFM models, each undergoing five recycling iterations. Following AFM structure generation, we applied AlphaFold-Multimer Local Interaction Score (AFM-LIS) and integrated Local Interaction Score (iLIS) to assess interaction confidence^32^. Positive PPIs were called using cutoffs (iLIS >= 0.223) derived from positive reference sets and matched control sets. The analysis code and cutoff details for iLIS analysis are available at https://github.com/flyark/AFM-LIS. Prediction details (Prediction scores, structures and PAE maps) are available in FlyPredictome (https://www.flyrnai.org/tools/fly_predictome/).

### Calcium Imaging Assays

Calcium mobilization in *Drosophila* S2R^+^ cells was measured using the fluorescent dye Fluo-4 AM (Thermo Fisher Scientific), adapted for live-cell imaging in ibidi µ-Slide 8 Well chambered coverslips (ibidi 80826-G500) pre-coated for optimal cell attachment. S2R^+^ cells stably expressing AkhR or CapaR were seeded in ibidi 8-well chamber slides at a density of 3×10D cells/well and allowed to attach overnight at 25°C. 5 µg Fluo-4 AM (ThermoFisher, F14201) was dissolved in 5 µL DMSO. From this, 2 µL of the stock solution was diluted into 405 µL of calcium-containing imaging buffer as mentioned in the previous study^33^ (final Fluo-4 AM concentration 10 µM). 20 µL of this working solution was added per well, followed by incubation for 1 hour at 22°C. After loading, cells were washed once with calcium-containing imaging buffer (composition as per ibidi slide instructions or standard Schneider’s-based buffer with Ca²^+^). Cells were maintained in the same buffer for imaging. Calcium flux was stimulated by addition of 10 µg/mL synthetic AKH peptide and imaging was performed on a spinning-disk confocal or epifluorescence microscope with 488 nm excitation and 520 nm emission. Time-lapse images were acquired at 1–2 second intervals immediately following AKH addition. Fluorescence was quantified using ImageJ by calculating ΔF/FD from regions of interest within individual cells, normalized to baseline (pre-stimulation) intensity. Experiments were conducted in triplicate, and mean calcium responses were averaged across >50 cells per condition.

### Morphological and Cytoskeletal Remodeling Assays

Morphological changes in S2R^+^ cells upon GPCR activation were assessed by both microscopic imaging and F-actin staining. For live-cell morphology observations, AkhR-expressing cells were plated on glass coverslips (coated with concanavalin A to promote adhesion) and treated with 10 µg/mL AKH peptide. Phase-contrast time-lapse microscopy was used to monitor cell shape changes. Within 1 minute of AKH stimulation, cells exhibited membrane ruffling and onset of cell spreading, which intensified over ∼3 minutes. To visualize dynamic actin remodeling in live cells, we utilized a cell-permeant F-actin probe (Invitrogen, A57243). Cells were stained as per reagents’ manual instructions and imaged live during AKH stimulation by spinning-disk confocal microscopy. This revealed rapid reorganization of the actin cytoskeleton in AkhR cells following AKH addition, an effect not seen in vehicle-treated controls and more pronounced than the response observed in CapaR cells. For fixed-cell analyses of F-actin, cells were subjected to prolonged agonist treatment to capture downstream cytoskeletal phenotypes. AkhR and CapaR stable cells were treated with AKH (10 µg/mL) or Capa-1 (10 µM) continuously for 48 hours to induce maximal cytoskeletal changes. Cells were then fixed in 4% paraformaldehyde in PBS for 15 minutes at room temperature and permeabilized with 0.1% Triton X-100 for 5 minutes. Actin filaments were stained with Alexa Fluor 568–phalloidin (Thermo Fisher, 1:200 dilution in PBS with 1% BSA) for 20 minutes. Nuclei were counterstained with DAPI (5 µg/mL). Coverslips were mounted in ProLong Gold antifade reagent (Invitrogen), and samples were imaged on a Zeiss LSM 880 confocal microscope with a 40× oil objective. In AkhR-expressing cells, prolonged AKH stimulation caused pronounced cell spreading with extensive cortical F-actin networks and stress fiber formation, whereas untreated cells or CapaR cells showed only mild changes. Quantitation of cell spreading was performed by measuring cell planar area and circularity from phalloidin-stained images using ImageJ. At least 50 cells per condition were analyzed in three independent experiments.

### Immunofluorescence and Receptor Localization

Immunofluorescence microscopy was used to examine the subcellular localization of AkhR and CapaR under various conditions. Stable AkhR or CapaR-expressing S2R^+^ cells were seeded on poly-lysine-coated coverslips and either left untreated or treated with AKH (10 µg/mL, 1 hour) to examine any ligand-induced internalization. Cells were fixed with 4% paraformaldehyde for 10 minutes and permeabilized with 0.1% Triton X-100 in PBS for 5 minutes. After blocking with 5% normal goat serum in PBST (PBS + 0.1% Tween-20), cells were incubated overnight at 4°C with primary antibodies against the receptors’ epitope tag (mouse anti-VHH05, 1:1000) to label AkhR or CapaR. The next day, samples were washed and incubated with Alexa Fluor 488–conjugated goat anti-mouse IgG secondary antibody (1:1000, Thermo Fisher) for 1 hour at room temperature.

For co-localization studies, cells were co-stained with organelle markers^34^. The following primary antibodies were used in combination with AkhR stainings: rabbit anti-Golgin245 (1:500, Golgi marker), rabbit anti-Calnexin99A (1:500, ER marker), and mouse anti-Hrs (1:200, early endosome marker). Appropriate Alexa Fluor 568– or 647–labeled secondary antibodies were applied. Lysosomes were labeled by adding LysoTracker™ Deep Red (Invitrogen, L12492) 50 nM to live cells for 5 minutes prior to fixation. After stainings, coverslips were mounted and confocal images acquired (W1 Yokogawa Spinning disk with 50 um pinhole disk). For each condition, Z-stack images were collected and maximum intensity projections analyzed. Pearson’s correlation coefficients for co-localization between AkhR and organelle markers were calculated using ImageJ. Representative images are shown with consistent contrast settings, and at least 10 fields per condition were examined.

### Metabolic Labeling and Click-Chemistry Detection of Palmitoylation

To directly assess protein palmitoylation, we performed metabolic labeling with an alkyne-palmitate analog followed by click chemistry detection. We focused on Gαq, the heterotrimeric G protein α-subunit, as it contains an N-terminal cysteine known to be palmitoylated. S2R^+^ cells (wild-type and mutant lines as specified) were transiently transfected with a plasmid encoding *Drosophila* Gαq with a C-terminal 3×FLAG tag. 24 hours after transfection, cells were incubated with 50 µM 17-octadecynoic acid (17-ODYA), an alkyne-functionalized palmitate analog (Cayman Chemical, Cas# 34450-18-5), added to the culture medium. Cells were metabolically labeled in the presence of 17-ODYA for 24 hours at 25°C. Control dishes without 17-ODYA were maintained in parallel. After labeling, cells were washed with PBS and lysed in RIPA buffer (50 mM Tris-HCl pH 7.5, 150 mM NaCl, 1% NP-40, 0.5% sodium deoxycholate, 0.1% SDS) supplemented with protease inhibitors. Cleared lysates were subjected to immunoprecipitation by incubating with anti-FLAG M2 agarose beads (Millipore-Sigma, A2220-5ML) for 3 hours at 4°C to pull down 3×FLAG–Gαq. Beads were washed three times with lysis buffer, and the bound proteins were then processed for click chemistry. The immunoprecipitated protein on beads was resuspended in click reaction buffer containing TAMRA-azide, 5-carboxytetramethylrhodamine-azide, 50 µM (Lumiprobe, D7130), CuSOD (1 mM), Tris(3-hydroxypropyltriazolylmethyl)amine (THPTA or TBTA) ligand (100 µM), and tris(2-carboxyethyl)phosphine, TCEP, 1 mM (Goldbio, 51805-45-9). The copper-catalyzed azide–alkyne cycloaddition (CuAAC) reaction was carried out at room temperature for 1 hour with gentle agitation. This reaction covalently attaches the TAMRA fluorophore to alkyne-labeled (palmitoylated) proteins. After the click reaction, samples were boiled in SDS sample buffer and resolved by SDS-PAGE. Subsequently, proteins were transferred to a PVDF membrane for Western blotting (described below) to detect total Gαq and detect TAMRA fluorescence (excitation/emission ∼555/580 nm).

To verify equal Gαq protein loading, the same gel was immunoblotted with anti-FLAG M2 antibody; this confirmed that differences in TAMRA signal were not due to different protein levels. Using this assay, we quantified palmitoylation levels by normalizing TAMRA fluorescence intensity to the anti-FLAG blot intensity for Gαq in each sample.

### Western Blotting

For protein analysis, cells were lysed in ice-cold lysis buffer appropriate to the experiment (RIPA buffer for most analyses, or a mild NP-40 buffer for membrane proteins: 20 mM Tris pH 7.5, 100 mM NaCl, 1% NP-40, 5% glycerol) with protease inhibitor cocktail (Roche). Lysates were clarified by centrifugation at 14,000×g for 15 min. Protein concentrations were determined by BCA assay (Pierce), and equal amounts of protein (typically 20–30 µg per lane) were mixed with 4× Laemmli sample buffer with β-mercaptoethanol, boiled for 5 min, and loaded on SDS-PAGE gels (4– 20% gradient). After electrophoresis, proteins were transferred to PVDF membranes (Millipore) using a semi-dry transfer apparatus. Membranes were blocked for 1 hour in TBST + 5% nonfat milk and probed with primary antibodies overnight at 4°C. The following primary antibodies were used for immunoblotting: mouse anti-VHH05 (1:1000, GenScript) to detect VHH05-tagged AkhR/CapaR, rabbit anti-AkhR (1:500, custom polyclonal against C-terminal peptide), mouse anti-FLAG M2 (1:1000, Sigma) for FLAG-tagged proteins, and mouse anti-Actin (1:2000, MP Biomedicals) as a loading control. After washing, membranes were incubated with horseradish peroxidase (HRP)-conjugated secondary antibodies (goat anti-mouse or anti-rabbit IgG, 1:5000, Jackson ImmunoResearch) for 1 hour at room temperature. Blots were developed with an ECL substrate (SuperSignal West Dura, Thermo) and imaged using a chemiluminescent detector (Azure Biosystems). Densitometry was performed using ImageJ; band intensities for target proteins were normalized to Actin. For example, in clonal AkhR-expressing cells, we observed that RNAi knockdown of *CG34449* (*Zdhhc8*) or *CG5447* (*Golgin A7*) led to a marked reduction in AkhR protein levels compared to controls. These blots confirmed that the hits from the screen are required to maintain steady-state AkhR protein abundance. Each Western blot experiment was repeated at least three times with independent biological replicates.

### RNAi Knockdown and CRISPR Validation of Screen Hits

To validate top hits from the CRISPR screen, we performed both RNA interference (RNAi) knockdown and individual gene knockout experiments in S2R^+^ cells. Details of RNAi sequence can be accessed at https://www.flyrnai.org/up-torr/ using Drosophila RNAi Screening Center IDs: DRSC32911 DRSC32910 DRSC39517 DRSC26407 DRSC35374 DRSC15801 DRSC03329 DRSC41888 and DRSC28694. For RNAi in S2R^+^ cells, double-stranded RNAs (dsRNAs) were synthesized by in vitro transcription (MEGAscript T7 kit, Thermo) from PCR templates containing T7 promoters at both ends. Templates targeting *CG34449 (Zdhhc8)* and *CG5447 (Golgin A7*) were designed using the DRSC dsRNA library resource, amplifying ∼500 bp gene-specific regions with no 21-nt off-target matches. Control dsRNAs targeting *Rho1* (an essential gene, positive control for assay) and *GFP* (negative control) were also generated. S2R^+^ cells stably expressing AkhR were plated in 6-well plates (1×10^6 cells/well) in serum-free Schneider’s medium and incubated with 10 µg of dsRNA per well. After 1 hour at 25°C, 2 mL of complete medium with serum was added to each well. Cells were incubated for 4 days to allow knockdown, with a half-media change on day 2. Knockdown efficiency was confirmed by qRT-PCR (for mRNA reduction) and by observing any expected phenotypes (e.g., *Rho1* dsRNA caused cell rounding within 4 days as a positive control). For the AkhR pathway, Western blotting of cell lysates after RNAi confirmed that *CG34449* and *CG5447* dsRNAs substantially reduced AkhR protein levels relative to *GFP* dsRNA controls. Functional assays (calcium imaging and morphology assays as described above) were performed on knockdown cells to ensure that the phenocopied effects were consistent with the CRISPR knockout lines.

For gene knockout validations, we generated clonal S2R^+^ cell lines with independent mutations in *CG34449* and *CG5447*. Using the stable Cas9-expressing S2R^+^ line, we transfected plasmids expressing single sgRNAs targeting either *CG34449* or *CG5447* (designed using the CRISPR Optimal Target Finder tool) along with a puromycin resistance marker. After 48 hours, puromycin (5 µg/mL) was applied for 5 days to enrich for transfected cells. Surviving cells were then plated by limiting dilution in 96-well plates to isolate clonal populations. Clones were screened by high-resolution melting analysis and Sanger sequencing of PCR-amplified target loci. Four independent knockout clones for each gene (KO1 and KO2) with frameshift indels in all alleles were expanded for analysis. For CG34449, the sgRNAs used were: CGGACTGACCGGCTTCCACA, CCAATGCAGTTGTTCACCCC, TGTGAATGGACAGTGAGACG, and CTATGTGAAGAGTCATCCAT. For CG5447, the sgRNAs used were: TCTTCGAGGCCACCATCAAT, CCTCCGTCAAGTTCCATACA, GCAGGATCCCTCCTCCGCCT, CAGCCGTGTATGGAACTTGA, and GCAGGATCCCTCCTCCGCCT. These clonal knockouts were verified by Western blot to completely lack ZDHHC8 or Golgin A7 protein (for Golgin A7, an HA-tagged version was transiently expressed to confirm antibody specificity). The *CG34449 (Zdhhc8)* and *CG5447 (Golgin A7*) knockout clones displayed no obvious growth defect in basal conditions but showed specific phenotypes upon AKH stimulation: they remained viable with minimal cytopathic effect even under prolonged AKH treatment, in stark contrast to wild-type cells. Knockout cells also failed to undergo the actin cytoskeletal collapse and cell shape changes seen in AKH-stimulated wild-type cultures. These observations mirrored the phenotype of *G*α*q*-knockout cells and supported that the loss of ZDHHC8 or Golgin A7 confers resistance to AKH-induced signaling (consistent with the screen results). Additionally, calcium imaging confirmed that AKH-induced Ca²^+^ flux was significantly blunted in the *CG34449* and *CG5447* knockout lines compared to control cells (comparable to the effect of *G*α*q* loss). Thus, both RNAi and CRISPR validation experiments substantiated the role of ZDHHC8 and Golgin A7 in AkhR/Gαq signaling.

### Flow Cytometry Analysis

Flow cytometry was used at multiple stages of this study: to monitor the composition of mixed cell populations during the CRISPR screen, to assess cell viability, and to quantify changes in cell size/granularity (FSC/SSC) associated with morphological remodeling. All flow cytometry was performed on a BD LSRFortessa analyzer equipped with 405 nm, 488 nm, and 561 nm lasers. For each sample, at least 20,000 events were recorded and analyzed using FlowJo v10 software.

Screening Co-culture Analysis: During the AKH selection screen, aliquots of the mixed GFP^+^ library and BFP^+^ *G*α*q*-knockout cell population were taken every 2 days to track their relative abundance. Cells were harvested from culture, washed in PBS, and stained with 1 µg/mL DAPI just prior to analysis (to discriminate live/dead cells). The flow cytometer was configured to detect GFP (488 nm excitation, 530/30 nm emission) and BFP (405 nm excitation, 450/50 nm emission) signals. Forward scatter (FSC) and side scatter (SSC) parameters were used to gate single cells and exclude debris. Live cells were gated as DAPI^⁻^ (excluding dead, permeable cells). Within the live-cell gate, the proportions of GFP^+^ and BFP^+^ cells were measured. We observed a steady decrease in the percentage of GFP^+^ cells and a corresponding enrichment of BFP^+^ cells in the AKH-treated population over time. By the end of the selection, GFP^+^ cells comprised <10% of the live population, compared to ∼50% in the untreated control, reflecting the selective loss of AKH-responsive (signaling intact) cells. These flow cytometry data guided the timing of our endpoint harvest for sequencing (ensuring sufficient depletion of sensitive cells).

Forward/Side Scatter for Morphological Complexity: We also employed flow cytometry to quantify changes in cell size and internal complexity resulting from receptor activation. AkhR-expressing S2R^+^ cells (and control S2R^+^ cells lacking AkhR) were treated ± AKH (10 µg/mL, 48 hours), then gently lifted from plates with cell dissociation buffer and analyzed for FSC and SSC distribution. We found that AKH treatment led to a pronounced increase in SSC values in AkhR-expressing cells compared to untreated cells. SSC reflects light scattering due to cellular granularity and internal complexity; the higher SSC in AKH-treated AkhR cells is consistent with extensive cytoskeletal reorganization and organelle redistribution in these cells. In contrast, control cells without AkhR showed no significant shift in SSC upon AKH exposure. Forward scatter (cell size) in AkhR cells was slightly increased or unchanged, indicating that the cells spread rather than shrank (spreading increases cross-sectional area but also increases internal complexity). These flow cytometry measurements of FSC/SSC provided a quantitative, population-level confirmation of the morphological changes observed microscopically. For each condition, median FSC and SSC values were recorded, and the relative change in SSC (ΔSSC) due to treatment was calculated. The ΔSSC for AKH-treated AkhR cells was ∼+30–40% above baseline, whereas ΔSSC for AKH-treated control cells was ∼0%, in line with visual observations.

All flow cytometry experiments included appropriate single-color controls and compensation for fluorescence overlap between GFP and BFP channels. Data were analyzed by first gating live single cells (FSC/SSC gating, DAPI exclusion), then gating on fluorescence channels as needed. Plots of SSC vs. counts were generated in FlowJo to compare distributions. Flow cytometry core facilities at Harvard Medical School were used for instrument access.

### Statistical Analyses

Data are presented as mean ± standard error of the mean (SEM) unless indicated otherwise. All experiments were repeated at least three times independently to ensure reproducibility. Statistical analyses were performed using GraphPad Prism 9. For comparisons between two groups, two-tailed unpaired Student’s *t*-tests were used. For multi-group comparisons, one-way or two-way analysis of variance (ANOVA) was performed, followed by Tukey’s or Dunnett’s post hoc tests to correct for multiple comparisons, as appropriate. The specific statistical test used for each experiment is indicated in the figure legends or results. Significance levels were set at *P* < 0.05. For the CRISPR screen data, MAGeCK analysis (as described above) provided false discovery rates (FDR) for each gene; hits were considered significant based on an FDR threshold of 5% (or 10% for suggestive hits). In graphs, asterisks denote statistical significance (*P* < 0.05, P < 0.01, *P < 0.001). All data met the assumptions of the statistical tests used (e.g., approximately normal distribution for *t*-tests/ANOVA; equal variance between groups was confirmed by Levene’s test or comparing standard deviations). No statistical methods were used to predetermine sample sizes, but our sample sizes are similar to those generally employed in the *Drosophila* cell signaling field. Data visualization and curve fitting (for calcium traces and time-course data) were done in Prism. No outliers were excluded unless pre-specified criteria (e.g., technical failures) were met. All analyses were performed with the experimenter blinded to genotype or treatment where feasible (particularly for imaging and fly assays).

Throughout the study, key resources were obtained from commercial or public repositories. Specifically, 17-ODYA, Fluo-4 AM, and TAMRA-azide were purchased from the suppliers indicated above, and their quality was verified via expected positive control reactions. All plasmids, fly strains, and reagents are available upon request.

---

**Supplementary Figure 1.**
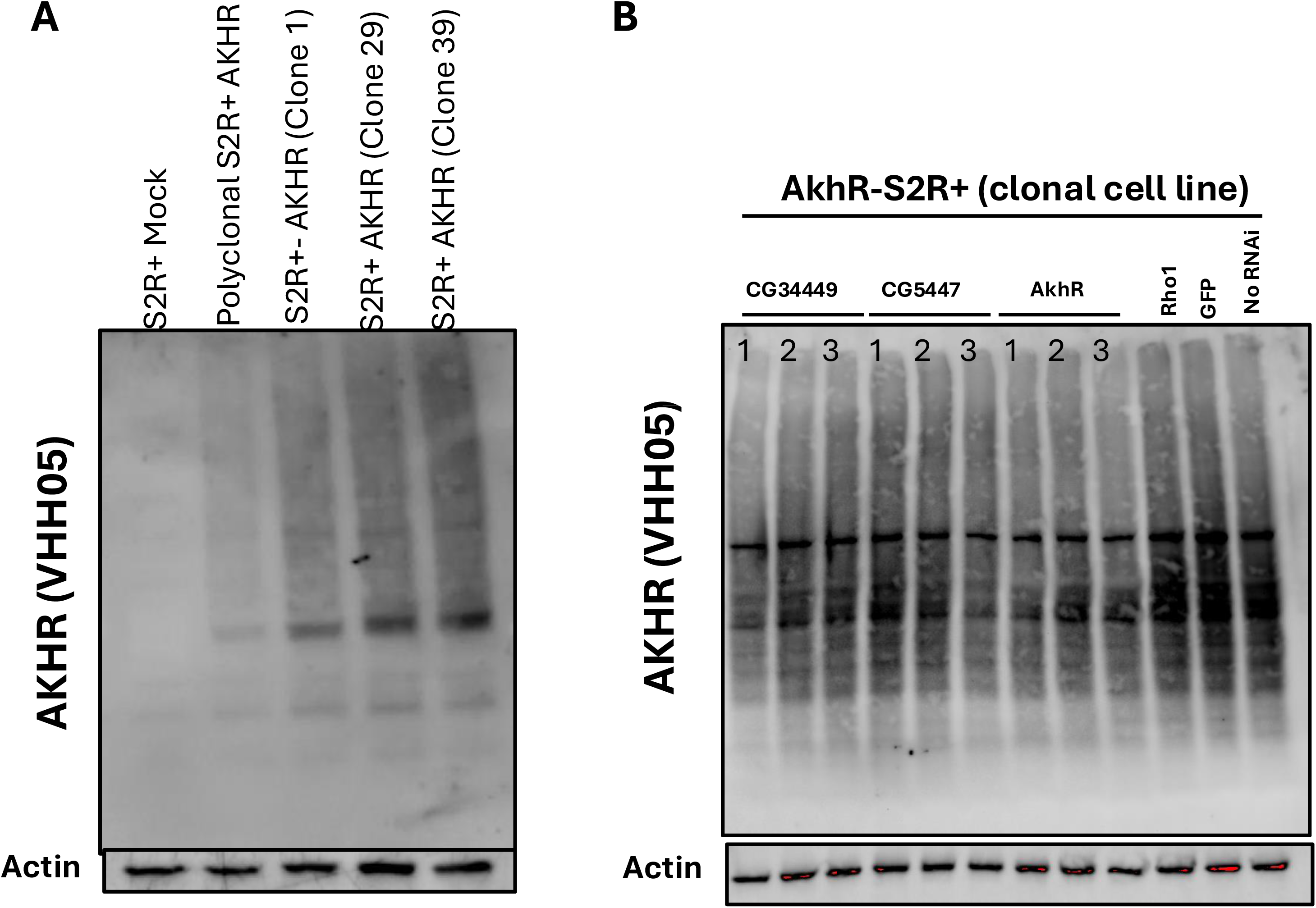
**(A)** Western blot analysis of multiple clonal S2R^+^ cell lines stably expressing VHH05-tagged AkhR. Clones 29 and 39 exhibit the highest levels of AkhR expression and were selected for all subsequent experiments. Mock-transfected and parental cells (S2R^+^ and S2R^+^-Cas9) show no detectable signal. **(B)** Western blot of AkhR protein levels in clone 39 S2R^+^-AkhR cells upon RNAi-mediated knockdown of screen hits *CG34449 (ZDHHC8), CG5447 (Golgin A7),* and *Akh*R itself. Three independent biological replicates are shown per condition. Knockdown of *CG34449* or *CG5447* reduces AkhR protein levels (similar to CRISPR KOs), phenocopying direct AkhR depletion. Actin serves as loading control.

**Supplementary Figure 2.**
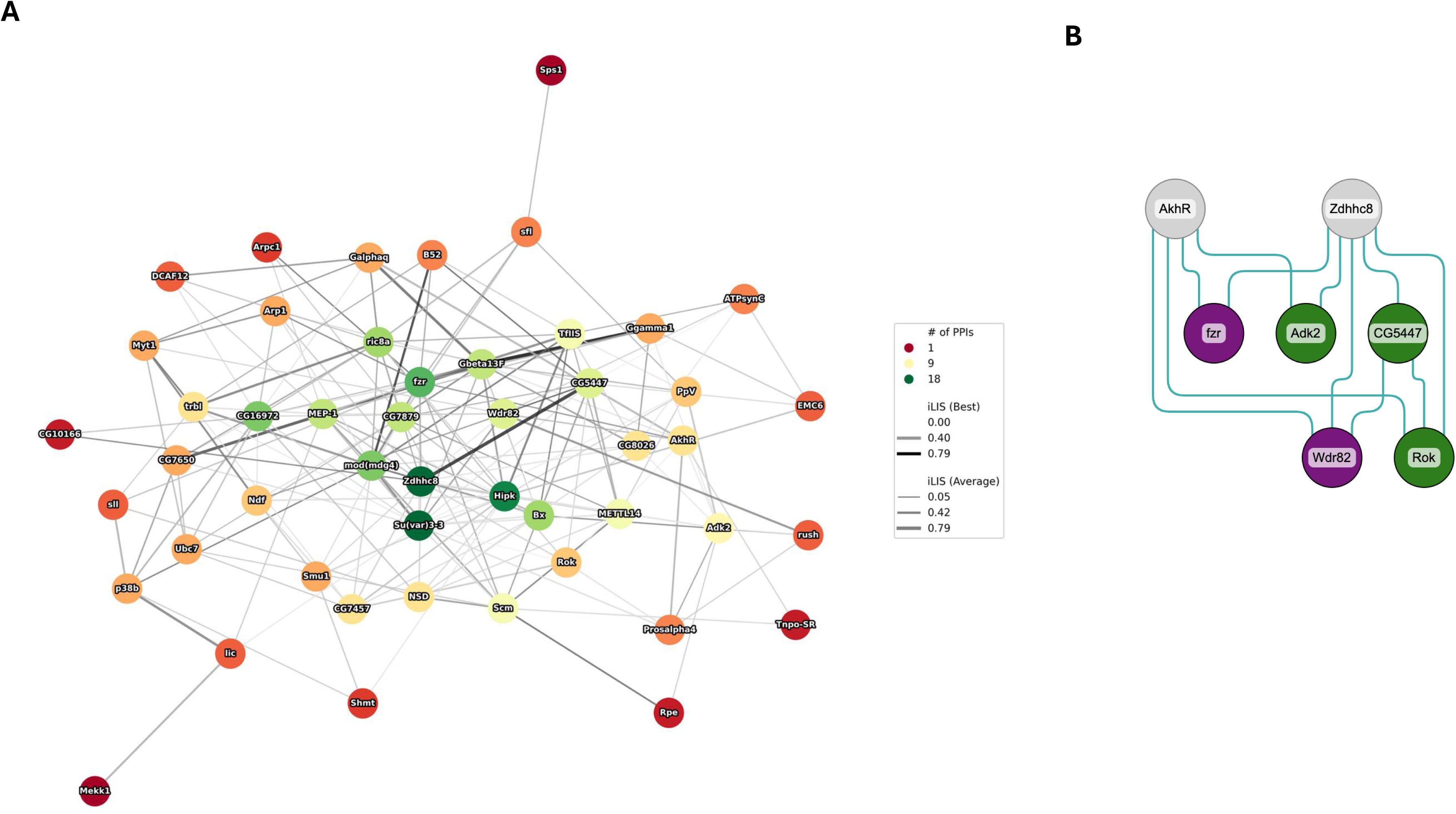
Network-level and pathway-focused visualization of predicted interactions among top CRISPR screen hits. **(A)** Network graph of AlphaFold-Multimer (AFM)-predicted protein-protein interactions (PPIs) among the top 50 genes identified in the AkhR-targeted CRISPR screen. Node color denotes the number of predicted PPIs (dark green = high connectivity; red = low), and edge thickness represents confidence of the predicted interaction (iLDDT interface score). Central clustering of proteins like ZDHHC8, CG5447, and Su(var)3–3 indicates potential roles as signaling hubs. **(B)** Schematic map of predicted PPIs linking key components of the AkhR–Gαq signaling axis. Core signaling elements (AkhR, ZDHHC8, CG5447) are connected to downstream modulators such as Rho kinase (Rok), Wek, Ear, and Akt2, suggesting that palmitoylation-dependent stabilization of receptor and G protein coordinates with cytoskeletal and survival pathways.

**Supplementary Figure 3:**
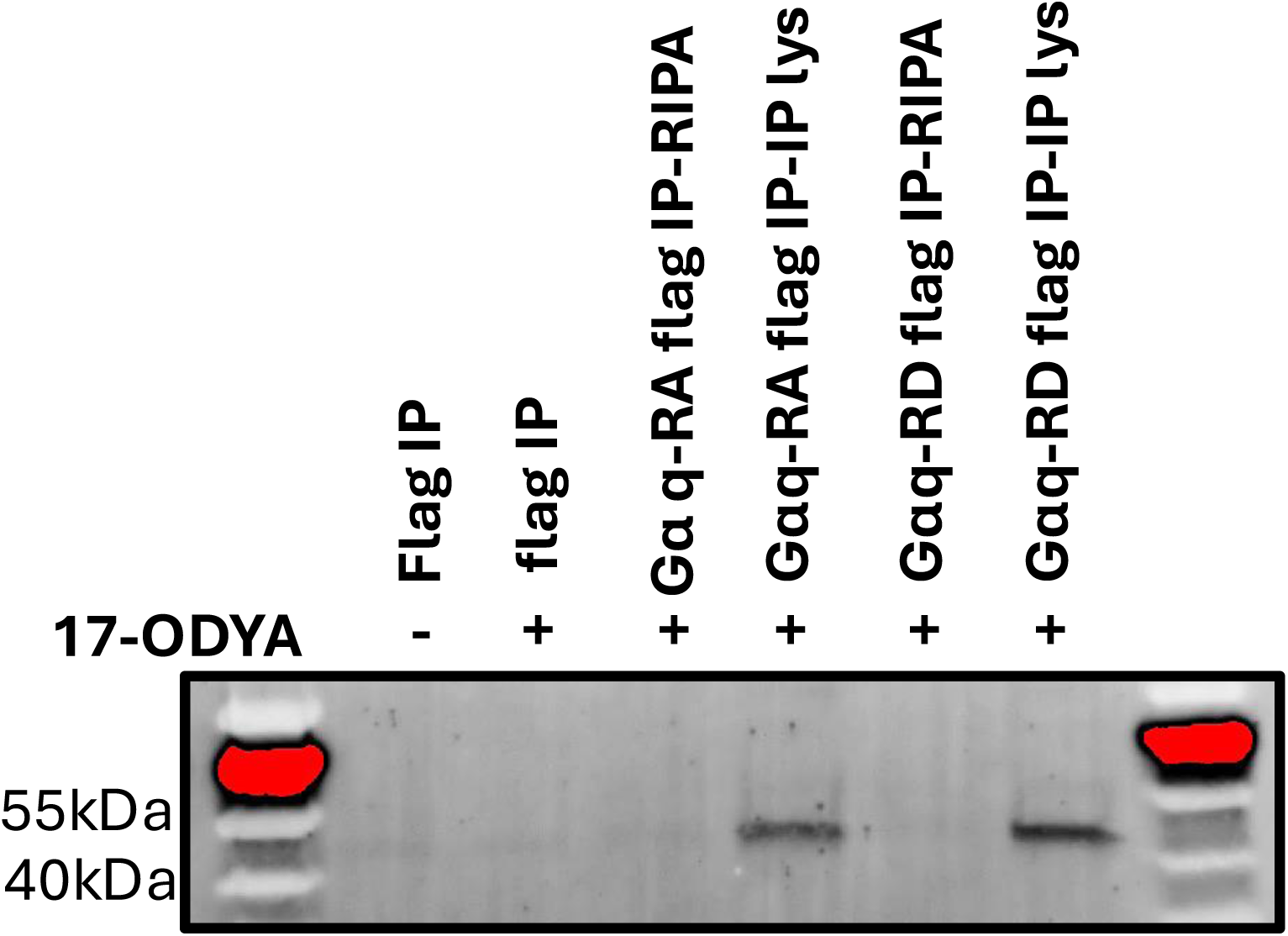
In-gel TAMRA fluorescence reveals specific labeling of G_α_q-3xFLAG following 17-ODYA incorporation. No signal is detected in the absence of 17-ODYA or when click chemistry reagents are omitted. Palmitoylation signal is detected in G_α_q-RA and G_α_q-RD variants, confirming that both incorporate 17-ODYA.

